# 3D dynamics of *trans* enhancer-promoter interactions in living *Drosophila embryos* reveals spatiotemporal thresholds for transcription activation

**DOI:** 10.1101/2025.03.17.643800

**Authors:** Hao Deng, Phillippe Valenti, François Payre, Bomyi Lim

## Abstract

While it is well acknowledged that specific enhancer-promoter interactions are essential for transcription, the spatiotemporal thresholds required for transcription initiation remain unclear. Here, we employed the phenomenon of transvection, where an enhancer activates the target gene in *trans* at the homologous position, to analyze the mechanism of long-range enhancer-promoter interactions. We combined the MS2- and PP7-based RNA labeling with the ParB/*parS* DNA labeling to simultaneously visualize active transcription and a target DNA locus in living *Drosophila* embryos. By quantifying the 3D enhancer-promoter distances in nuclei that exhibited active or inactive transcription, we found that sustained proximity between the enhancer and promoter is necessary and sufficient to initiate transcription. The enhancer-promoter proximity was established before transcription initiation and remained stable after the termination, suggesting the localization of an enhancer and the target promoters in a “transcription hub.” When transvection occurred, the *cis*-linked reporter gene showed a delayed activation and reduced mRNA production, suggesting homologous promoter competition in the hub. Our DNA-labeled co-transvection assay sheds insights into the activation criteria for the long-range enhancer-promoter interactions and provides a stepping stone for future endeavors to understand the complex gene regulation in 3D space.

## Introduction

The normal development of an organism relies on the precise regulation of genetic information and subsequent protein expression. While it is well known that mutations in protein coding regions result in significant phenotypic alterations, recent advancements in genome-wide studies have found that mutations in non-coding DNA regions can also cause disruptions in cellular functions, including human diseases, as well as play an essential role in evolution^1-6^. These regions often correspond to enhancer sequences that interact with the target promoters to initiate and regulate transcription^7^. Enhancers contain a specific combination of transcription factor binding sites, which determines the spatial and temporal dynamics of the target gene expression^8,9^. Notably, the regulatory control executed by enhancers is not dictated by their position relative to the target genes^10-12^. Enhancers can be located at varying distances from their target genes, ranging from immediately adjacent positions to several million base pairs away^13-17^. They may also be positioned upstream or downstream of the genes they regulate^10-12,16^. In *Drosophila,* the average genomic distance between an enhancer and its target promoter is around 10 kb^13^; in mammalian cells, these distances can range from tens to hundreds of kb^15^. In the case of a distal enhancer-promoter (E-P) pair, the presence of additional E-P pairs, transcription units, and regulatory elements in between can create a complex genomic environment, complicating the regulation of E-P interactions. Despite the long E-P genomic distance and complex genomic contexts, E-P interactions are highly specific^16^. Such precision is achieved by insulator proteins, which are recruited to their DNA binding sites, dimerize to induce proximity between enhancers and promoters, and, as a result, facilitate long-range E-P interactions^18^. For example, the gene *Sex combs reduced* (*Scr*) and its T1 enhancer are 25 kb apart, with *fushitarazu (ftz*) and its enhancer located in the intermediary genomic region. Through insulator-mediated pairing, the T1 enhancer interacts with its distant target gene *Scr*, bypassing the nearby *ftz* gene^19^.

Since the discovery of enhancers in the 1980s, the mechanisms underlying precise E-P interactions have been extensively studied^20-22^. Significant phenotypic changes can result from mutations in enhancers or promoters and disruptions in E-P communications^5,6^. However, studies on the conditions of E-P interactions that are required to initiate transcription have yielded contradictory findings. In *Drosophila* embryos, it was shown that enhancers and their target promoters need to be in physical proximity of around 350 nm to trigger transcriptional initiation^23^. On the other hand, studies in mammalian cells demonstrated no correlation – even a negative correlation – between E-P distance and transcriptional activity^24-29^. Despite these findings, the spatiotemporal characterization of long-range E-P interactions and their roles in transcriptional regulation remain largely unexplored. For instance, what are the distance and time criteria for distant enhancers and their target promoters to initiate transcription? Does the E-P interaction change after transcription termination? When a single enhancer interacts with multiple promoters, how does it affect the transcriptional activity of each promoter?

A live imaging-based co-transvection assay was used to characterize the dynamics of long-range E-P interactions in living *Drosophila* embryos with a high spatiotemporal resolution^30^. First discovered in 1954, transvection refers to a phenomenon where, in diploid cells, an enhancer activates its target promoter located on the paired homolog in *trans* with the help of insulator complexes that facilitate chromosome pairing^31,32^. For example, two loss of function mutations of the same gene, one affecting *cis*-regulatory regions and the other protein coding sequences, can complement each other when placed in a *trans-*allelic combination; the intact enhancer acting in *trans* to activate the homologous allele that expresses wild type protein^33^. Since *trans*-activation requires an enhancer to interact with the target promoter on the homologous allele over a long distance, transvection is well suited to examine the dynamics of long-range E-P interactions. Implementing live imaging techniques to study the transvection phenomenon, a previous study demonstrated that a single enhancer can simultaneously co-activate both homologous promoters in *cis* and *trans*^30^. Additionally, the study reported that both homologous alleles are in proximity during *trans*-activation events^30^. However, this assay only visualized E-P interactions when both promoters are actively transcribed. Consequently, it couldn’t capture the spatiotemporal dynamics of the enhancer and promoters before and after transcription – information crucial for understanding the mechanisms of enhancer-mediated transcriptional regulation.

To improve the assay and visualize E-P interactions even when transcription is inactive, we implemented a live DNA labeling technique to establish a DNA-labeled co-transvection assay. We quantified the three-dimensional (3D) E-P distance throughout the nuclear cycle 14 (NC14) (∼40 min) and demonstrated that enhancers and their target promoters should sustain physical proximity for at least a few minutes to initiate transcription. We also presented evidence that enhancers and their target promoters established a stable association before transcriptional initiation and remained in proximity after transcription was terminated. Analysis of transcriptional dynamics showed that activation of the *trans-*linked reporter gene led to delays in transcriptional initiation and a reduction in mRNA production of the *cis-*linked reporter gene, suggesting a competitive relationship between the homologous promoters when regulated by a shared enhancer. Together, our data support the hypothesis that transcription co-activation occurs in a stable transcription hub where enhancers and promoters are in close proximity, likely improving access to a reservoir of limiting transcriptional machinery.

## Results

### DNA-labeled co-transvection assay visualizes long-range enhancer-promoter interactions

The co-transvection assay combines the transvection model with nascent RNA live imaging to investigate the dynamics of E-P interactions^30^. The assay contains a pair of *gypsy* insulators, the *snail* distal enhancer with a *cis-*linked reporter gene, and a *trans-* linked reporter gene at the homologous positions (Fig. 1a). The *trans*-linked reporter gene can be activated only when the enhancer on the homologous allele interacts with the promoter in *trans*. The *trans*-activation is facilitated by the binding of insulator proteins to the 12 Suppressor of Hairy-wing (Su(Hw)) binding sites in the *gypsy* insulators^34^. The binding and dimerization of insulator proteins stabilize homolog pairing such that transcription in *trans* would be absent if either or both *gypsy* insulators at the homologous positions are missing^30,35^. With the placement of the enhancer and promoter on separate alleles, the co-transvection assay allows visualization of the long-range E-P communications.

**Figure 1.**
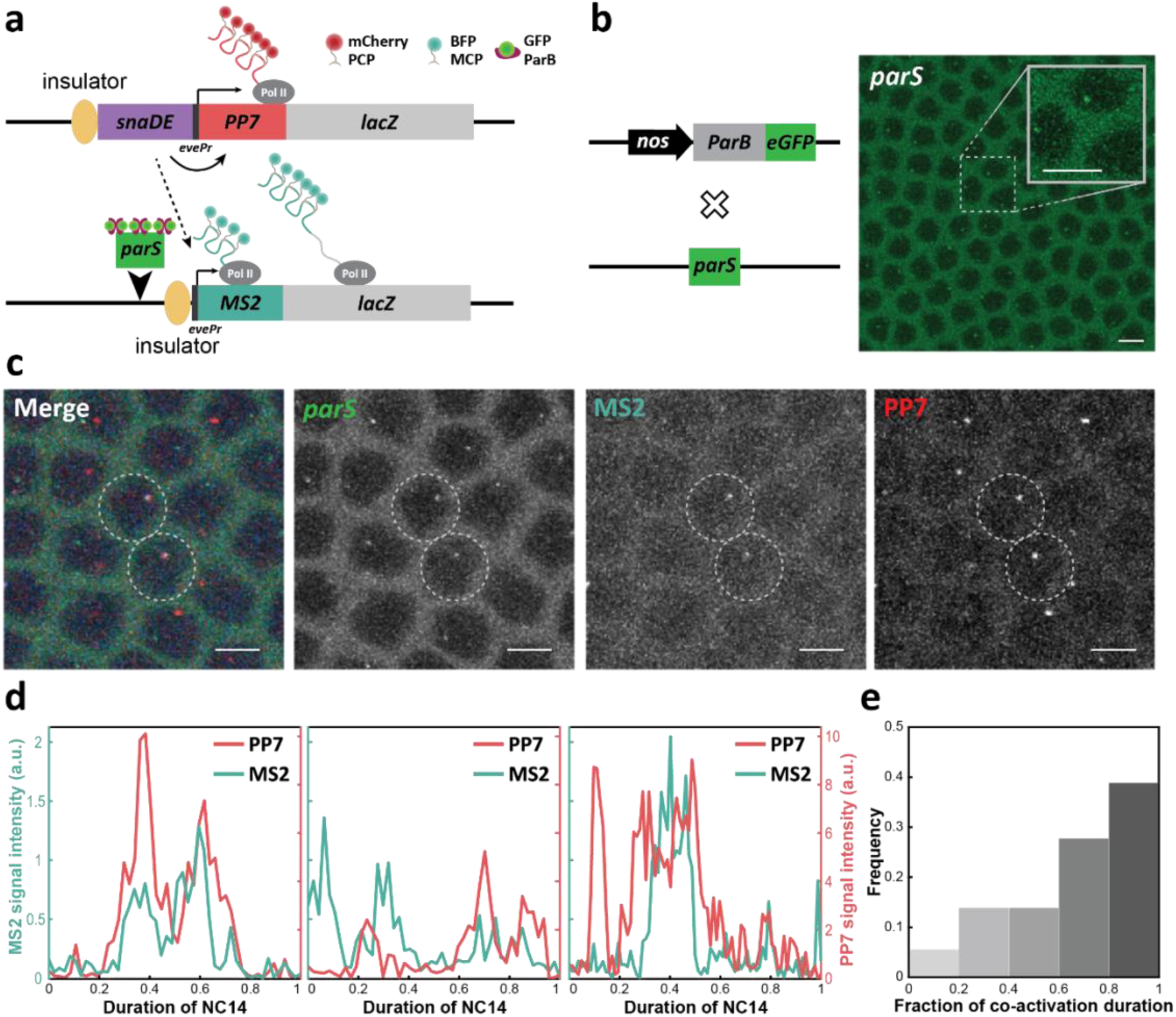
The DNA-labeled co-transvection assay to visualize long-range E-P interactions. **A)** Schematic of the DNA-labeled co-transvection assay. The co-transvection assay contains a pair of *gypsy* insulators, the *snail* distal (snaDE) enhancer with a *cis-*linked reporter gene, and a *trans-*linked reporter gene at the homologous positions. The *parS* sequences are inserted directly upstream of the *gypsy* insulator of the enhancer-less allele. PCP::mKate, MCP::BFP, and ParB::eGFP bind to *PP7* stem loops, *MS2* stem loops, and the *parS* DNA sequence, respectively. The snaDE enhancer regulates both *PP7* and *MS2* reporter genes, while it interacts more frequently with the *cis-*linked *PP7* reporter gene (solid arrow) than the *trans-*linked *MS2* reporter gene (dashed arrow). **B)** Schematic of the ParB/*parS* DNA labeling system where the *parS* sequence is inserted into the DNA, and ParB::eGFP is deposited maternally through the *nanos* promoter (left). Live image snapshot of an embryo showing the ParB::eGFP signal bound to the *parS* sequence (right). Since ParB directly binds to the DNA, the fluorescent signals should be active in all nuclei at all times. Scale bars, 5 μm. **C)** Snapshot of an embryo containing the DNA-labeled co-transvection assay shown in (A). The two white dashed circles highlight two nuclei with active transvection. Without *trans*-activation, only the PP7 and *parS* signals are active, each of which indicates the location of the enhancer and promoter, respectively. Scale bars, 5 μm. **D)** Representative MS2 and PP7 signal trajectories of individual transvecting nuclei. While the activation time of *cis-*linked *PP7* and *trans-*linked *MS2* reporter genes can vary from nuclei to nuclei, all transvecting nuclei show co-activation of PP7 and MS2. **E)** Relative frequency distribution of the fraction of MS2 and PP7 co-activated time frames over the duration of MS2 activity. When the *trans-*linked *MS2* reporter gene is activated, the *cis-*linked *PP7* reporter gene is most likely activated as well.

The MS2 and PP7 live imaging method was used to visualize the enhancer-containing allele and enhancer-less allele. The dynamics of the reporter genes can be traced by inserting 24 copies of PP7 and MS2 DNA sequences into the 5’UTR of the *cis-* and *trans-*linked *lacZ* reporter genes, respectively (Fig. 1a). Upon transcription, PP7 and MS2 sequences form mRNA stem loops, each bound by two copies of maternally deposited PP7 coat proteins (PCP) or MS2 coat proteins (MCP) fused with fluorescent proteins^36-38^. When nascent transcripts are generated, corresponding fluorescent signals are instantaneously captured and represent the location and transcriptional activity of the reporter gene. In the co-transvection assay, the *trans-*linked reporter gene is tagged with MS2, such that the detection of MS2 signals corresponds to instances of transvection. The E-P distance can be approximated with the coordinates of PP7 and MS2 signals. However, this co-transvection assay visualizes the E-P interactions only when the enhancer co-activates both the *cis*- (*PP7*) and *trans*-linked (*MS2*) reporter genes. Given that the *trans-*activation only occurs in about 3-5% of nuclei and for a short time window during NC14 (Fig. 1c), the ability to trace the spatiotemporal dynamics of E-P interactions is limited.

To improve the tracking of E-P interactions throughout NC14 and overcome the challenge of visualizing the enhancer-less allele in the absence of *trans*-activation, we implemented the ParB/*parS*-mediated DNA labeling system. The ParB fused with fluorescent proteins bind and accumulate at the *parS* DNA sequences, allowing the visualization of a specific locus independent of transcriptional activation (Fig. 1b). We modified the co-transvection assay by inserting the *parS* sequences adjacent to the insulator on the enhancer-less allele that contains the *trans-*linked *MS2* reporter gene (Fig. 1a). MCP::BFP, mKate::PCP, and ParB::eGFP are maternally provided to detect MS2, PP7, and *parS*, respectively. The *cis-*linked *PP7* reporter gene has strong and continuous baseline activity, thus the positional information of the PP7 signal can function as a proxy for the enhancer position. Meanwhile, the coordinates of the *parS* or MS2 signal in the *trans* position can approximate the location of the promoter. In addition, MS2 signal is used to visualize transvection, by detecting transcriptional activation of the *trans-*linked *MS2* reporter gene. This DNA-labeled co-transvection live imaging assay allows simultaneous tracking of the long-range E-P interactions and subsequent transcriptional activity of both reporter genes throughout NC14.

We first examined the transcriptional activity of both *cis-* and *trans-*linked reporter genes regulated by a shared enhancer. Some nuclei showed activation of the *trans-*linked reporter gene, signaling the occurrence of *trans* E-P interactions. Interestingly, all *trans-* activation of the *MS2* reporter gene was accompanied by the activation of the *cis-*linked *PP7* reporter gene, exhibiting a time window of simultaneous PP7 and MS2 activation (Fig. 1d). In over 66% of *trans*-activating nuclei, more than half of the time frames when *trans-*activation occurred were accompanied by *cis*-activation (Fig. 1e). Additionally, 24% of *trans*-activating nuclei exhibited *trans*-activation (*MS2*) before the *cis-*activation (*PP7*) (Fig. 1d, center) despite the *cis-*linked reporter gene being physically proximal to the shared enhancer (Supplementary Fig. 1). This implies that the dimerization of insulators and long-range homolog pairing may occur in early NC14 to facilitate *trans-* activation. The varying activation timings of reporter genes illustrate how dynamic the transcription process is and highlight the importance of temporal information in understanding the mechanisms underlying transcriptional initiation.

### Enhancer-promoter proximity is required for transvection

We used our DNA-labeled co-transvection assay to further probe the spatial and temporal threshold of long-range E-P interactions required for transcription initiation. We defined “transvecting” nuclei (Transvection ON) as nuclei that exhibited active MS2 signals driven by the *trans*-linked promoter, and “non-transvecting” nuclei (Transvection OFF) as those with active PP7 signals driven by the *cis-*linked promoter with no *MS2* transcription (*trans*-activation) (Fig. 2a). The x-, y-, and z-coordinates of each fluorescent signal (MS2, PP7, or *parS*) were used to calculate the 3D distance between the *snail* enhancer and the *trans*-linked promoter. Transvecting nuclei exhibited an average E-P distance of around 506 ± 317 nm (Fig. 2b, center) during NC14 and 407 ± 258 nm (Fig. 2b, left) when transvection was active. These were much shorter than the E-P distance in non-transvecting nuclei, which showed a wide range of values with an average of around 3.30 ± 1.26 μm (Fig. 2b, right). Most transvecting nuclei demonstrated spatial proximity between the enhancer and promoter during NC14. Based on this observation, we conclude that enhancer-promoter proximity is a prerequisite for initiating transcription. In essence, an enhancer is unable to activate its target gene in *trans* without achieving this spatial closeness to the promoter.

**Figure 2.**
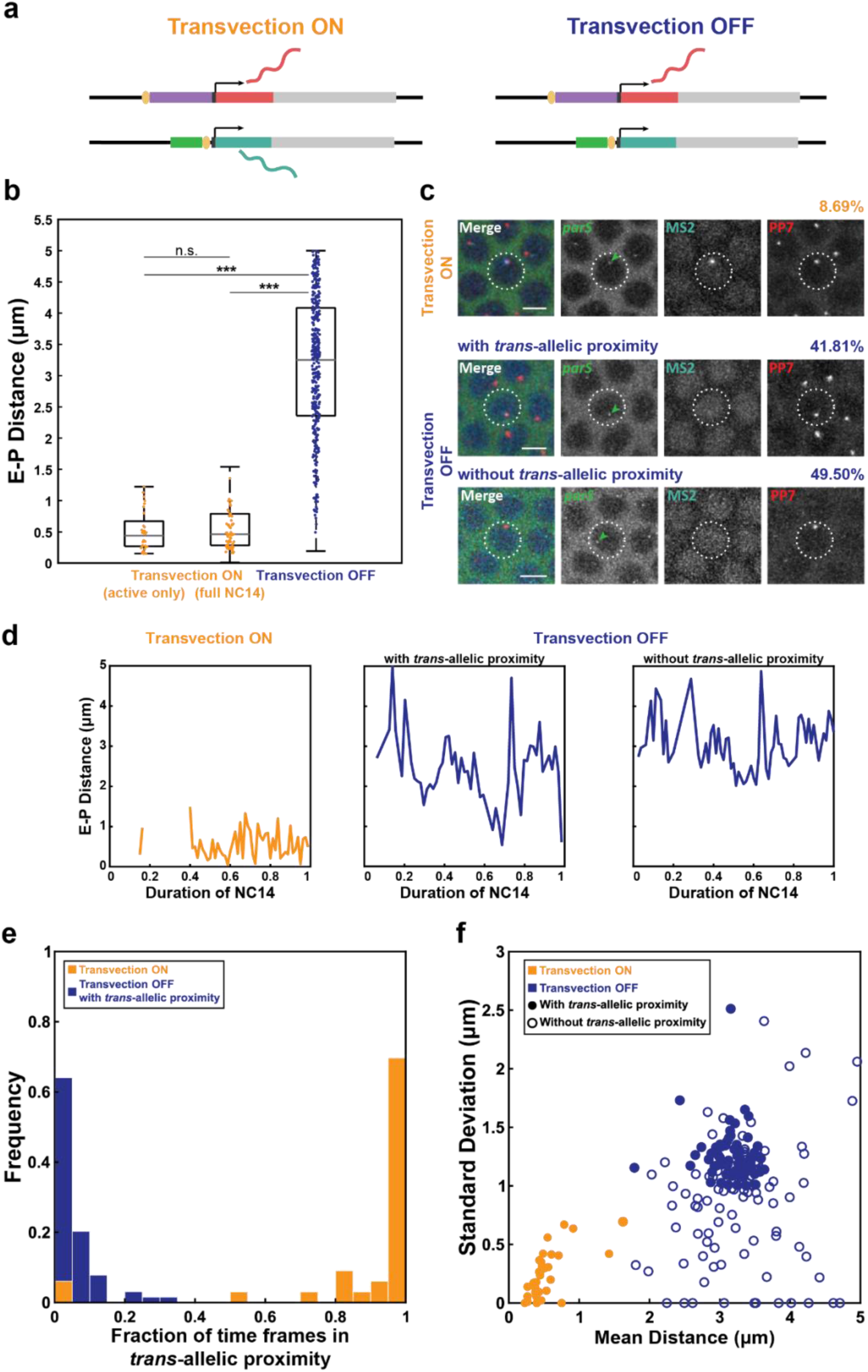
A sustained *trans*-allelic proximity is necessary and sufficient for transvection initiation. **A)** Simplified schematics of the DNA-labeled co-transvection assay showing “Transvection ON” and “Transvection OFF” status. **B)** Boxplots showing the 3D E-P distance in Transvection ON nuclei during active transvection, Transvection ON nuclei throughout NC14, and Transvection OFF nuclei in NC14. Each data point represents the E-P distance at a single time frame. In each box plot, the central gray line indicates the median, and the top and bottom edges indicate the 25th and 75th percentiles, respectively. The whiskers indicate the maximum and minimum of the dataset excluding outliers. The sample size of each box plot is 129, 180, and 3410, respectively, from 14 biological replicate embryos. n.s. indicates no statistical significance and ***p<1E-4 using the Welch’s *t*-test. **C)** Snapshots of Transvection ON nuclei (top), Transvection OFF nuclei with (center) and without (bottom) *trans-*allelic proximity. White dashed circles highlight nuclei representing each of the above cases. *parS*, MS2, and PP7 signals all overlap in Transvection ON nuclei. In Transvection OFF nuclei, no MS2 signal is observed, and only the nuclei with *trans*-allelic proximity show overlapping *parS* and PP7 signals. Scale bars, 5 μm. **D)** Representative E-P distance trajectories of Transvection ON nuclei (left), Transvection OFF nuclei with (center) and without (right) *trans-*allelic proximity. **E)** The relative frequency distribution of the fraction of time frames when the enhancer and the target promoter on the enhancer-less allele are in *trans*-allelic proximity. The *trans-*allelic proximity is transient in Transvection OFF nuclei (blue), while it is persistent in Transvection ON nuclei (orange). **F)** The scatter plot of means and standard deviations of E-P distance trajectories. Transvection ON nuclei (orange) are low in both average and standard deviation. Transvection OFF nuclei (blue) show high averages and a wide range of standard deviations, indicating that transient *trans-*allelic proximity is not sufficient for transcription initiation. 33, 63, and 104 Transvection ON, Transvection OFF with *trans*-allelic proximity, and Transvection OFF without *trans*-allelic proximity nuclei were analyzed from 14 biological replicates.

### Sustained enhancer-promoter association is needed to initiate transcription

While most transvecting nuclei exhibited E-P proximity, some non-transvecting nuclei also showed E-P proximity at some time frames (Fig. 2b and c). This observation raised the question of whether E-P proximity is sufficient to initiate transcription. We analyzed and compared the dynamics of the E-P distance between transvecting and non-transvecting nuclei over a full NC14 time course. We defined “*trans*-allelic proximity” as the state when the enhancer and promoter were less than 1 μm from each other. Surprisingly, we found that about half of the total nuclei exhibited *trans*-allelic proximity regardless of the transvection state (ON or OFF) (Fig. 2c and d). In fact, of these nuclei with proximity, only 17% demonstrated *trans*-activation of the *MS2* reporter (Fig. 2c). We plotted the relative frequency distribution of the fraction of time frames when the enhancer and promoter were in *trans-*allelic proximity in transvecting versus non-transvecting nuclei (Fig. 2e). We found that the proximity occurred transiently in non-transvecting nuclei, only lasting less than 5% of the total recorded time frames. On the other hand, the *trans*-allelic proximity associated with transvecting nuclei was much more stable, lasting over 90% of the total recorded time frames. Our findings suggest that proximity alone is insufficient to initiate transcription, and that duration and stability of the *trans*-allelic proximity state may play a critical role in transcriptional initiation.

To further probe the importance of the duration and stability of *trans-*allelic proximity in regulating transcription initiation, we analyzed the distance trajectories of transvecting and non-transvecting nuclei. The transvecting nuclei showed little change in E-P distance over time, indicating that stable E-P association is maintained throughout NC14 (Fig. 2d, left). In contrast, some non-transvecting nuclei exhibited consistently long E-P distances over time while others had stochastic and fluctuating distance trajectories, where the E-P distances changed significantly in a short time window (Fig. 2d, center and right). Such E-P distance fluctuations in non-transvecting nuclei can be interpreted as the free diffusion of two alleles when insulators fail to establish homolog pairing. To better characterize the stochastic nature of distance trajectories, we calculated the mean and standard deviation of the distance trajectories of transvecting and non-transvecting nuclei. In the scatter plot, transvecting and non-transvecting nuclei were clustered into two subgroups (Fig. 2f). As expected, in transvecting nuclei (Fig. 2f, orange-filled dots), the enhancer and promoter sustained stable association, demonstrated by their low average distances and standard deviations. On the other hand, the distance trajectories of non-transvecting nuclei (Fig. 2f, blue-filled and hollow dots) showed a wide range of standard deviations with high average E-P distances, which suggests the absence of stable allelic association. Taken a closer look, in non-transvecting nuclei that exhibited *trans-*allelic proximity (Fig. 2f, blue-filled dots), the enhancer and promoter diffused stochastically with no stable association despite the presence of proximity at some time frames in their distance trajectories. From the stochastic nature of the distance trajectories of non-transvecting nuclei, we conclude that transient *trans*-allelic proximity is not sufficient to trigger transcription despite its high probability of occurrence. In other words, stable E-P association is necessary for transcription initiation.

### Stable enhancer-promoter association is maintained before and after transvection

To determine the temporal threshold of E-P interactions needed for transcription initiation, we quantified the changes in the E-P distance of transvecting nuclei before and after transvection throughout NC14 (Fig. 3a). Before transvection, the distance trajectory showed that the E-P distance gradually decreased and reached a plateau at ∼500 nm around 4 minutes before the onset of transvection (Fig. 3b). After the termination of transvection, the E-P distance remained constant for around eleven minutes and gradually increased afterward (Fig. 3b), indicating the stable E-P interactions last longer than the duration of transcription.

**Figure 3.**
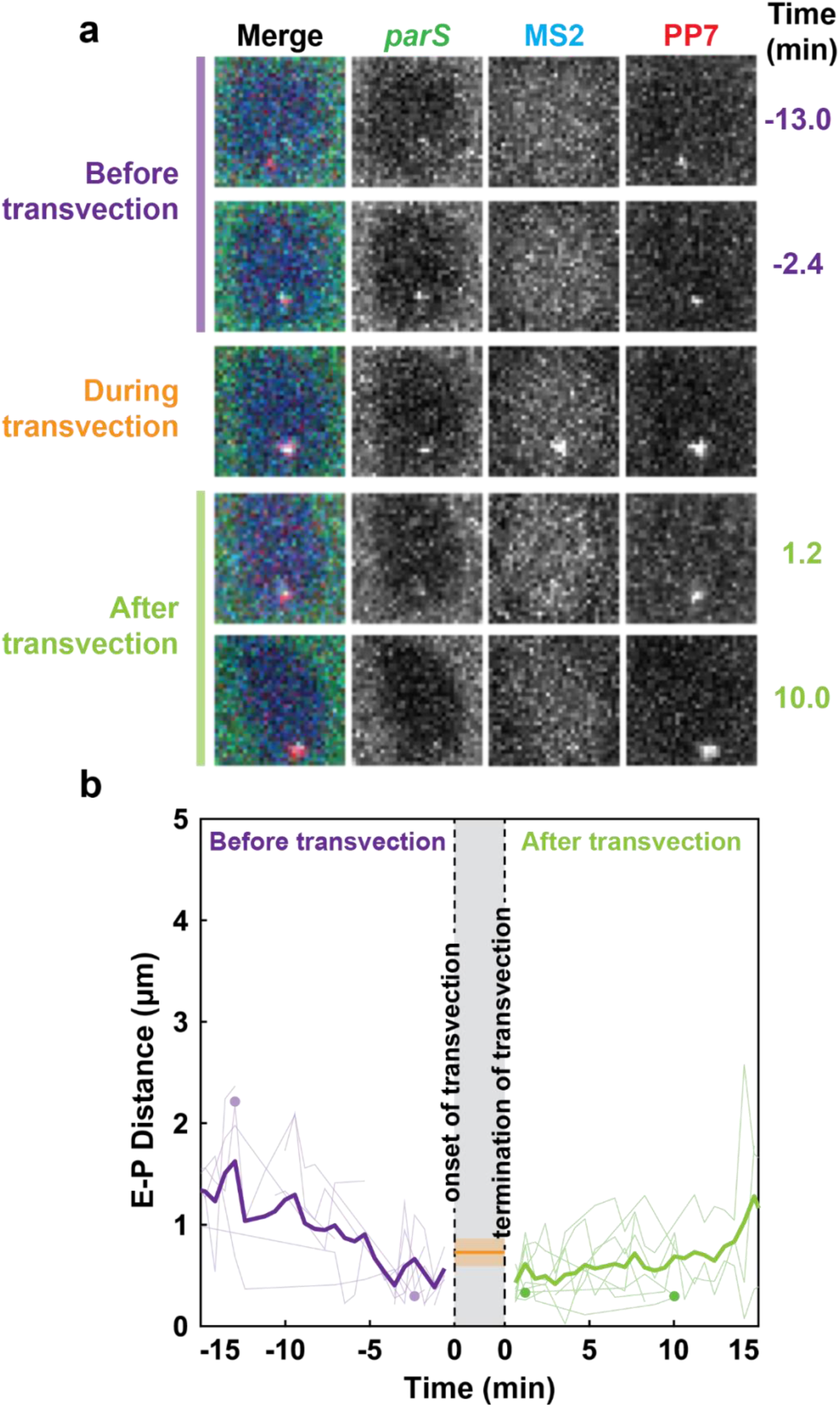
*Trans-*allelic proximity is observed before, during, and after active transvection. **A)** Snapshots of a nucleus at different time frames before, during, and after transvection. **B)** The E-P distance trajectories of Transvection ON nuclei before and after transvection. The “0” at the x-axis encompasses the duration of transvection from the onset to the termination of transvection. The light-colored trajectories are the raw E-P distance trajectories from each Transvection ON nucleus. The dark-colored trajectories are the average E-P distance trajectories. The circles correspond to the time points of the nucleus before or after transvection shown in Figure 3A. The line and shaded area of the orange band in the gray region indicates the median and range of the average E-P distance of each nucleus during transvection. The E-P distance gradually decreases and reaches a plateau around four minutes before the onset of transvection. The E-P distance remains constant and gradually increases around ten minutes after the termination of transvection. 12 and 10 Transvection ON nuclei in 9 and 7 biological replicates were analyzed for before and after transvection E-P distance analysis, respectively. The average E-P distance during transvection was calculated from 36 Transvection ON nuclei in 10 biological replicate embryos.

To test whether the E-P interaction criteria for transcription initiation is specific to the *gypsy* insulators, we replaced the *gypsy* insulators with the *homie* insulators in the DNA-labeled co-transvection construct (Supplementary Fig. 1a). Endogenously located near the *even-skipped* gene in *Drosophila*, *homie* contains binding sites for both Su(Hw) and CTCF insulator binding proteins^39^. Spatiotemporal conditions necessary for transcription initiation were similar between the *gypsy-* and *homie-*containing constructs, requiring a stable E-P proximity for a few minutes (Supplementary Fig. 2b and c). Moreover, while the average E-P distance of transvecting nuclei remained proximal throughout NC14, E-P distance gradually decreased to <500 nm around 4 minutes before the initiation of transvection, agreeing with the results from the *gypsy-*containing constructs (Supplementary Fig. 2d and e). Although *homie* insulators are not as efficient in homology pairing and facilitating transvection^30^, the initiation thresholds seem to be maintained as long as the homolog pairing can be established, regardless of the insulator strength.

The results from both *gypsy* and *homie* constructs support the “transcription hub” hypothesis and insinuate that a specific E-P interaction is established before transcriptional initiation, via homolog pairing of the insulators to prepare for transcription. As a model of enhancer-promoter interactions, the “transcription hub” hypothesizes that enhancers and promoters interact in a cluster of transcriptional machinery, as opposed to the classical looping model^30,40-42^. Our observations showed that the transcription hub seems to be maintained beyond the duration of transcription and is slowly dissociated. Therefore, such transcription hubs are likely a prerequisite for transcription and do not result from a mere and transient accumulation of the transcriptional machinery.

### Homologous promoter competition can cause delays in enhancer-promoter interactions

We conducted an in-depth analysis of transcription hub characteristics by examining the transcriptional dynamics of both *cis*-linked *PP7* and *trans*-linked *MS2* reporter genes (Fig. 4a). In transvecting nuclei, the *cis*-linked *PP7* reporter gene was activated at a faster rate compared to the *trans-*linked *MS2* reporter gene (Fig. 4b and c). This phenomenon could be due to the distance advantage of the *cis*-linked reporter gene, which is physically closer to the enhancer and does not require homolog pairing as the *trans*-linked promoter does. These results suggest that the enhancer and *trans-*linked promoter often need more time in homolog pairing to induce *trans*-activation while the enhancer is readily available to initiate transcription at the *cis*-linked promoter.

**Figure 4.**
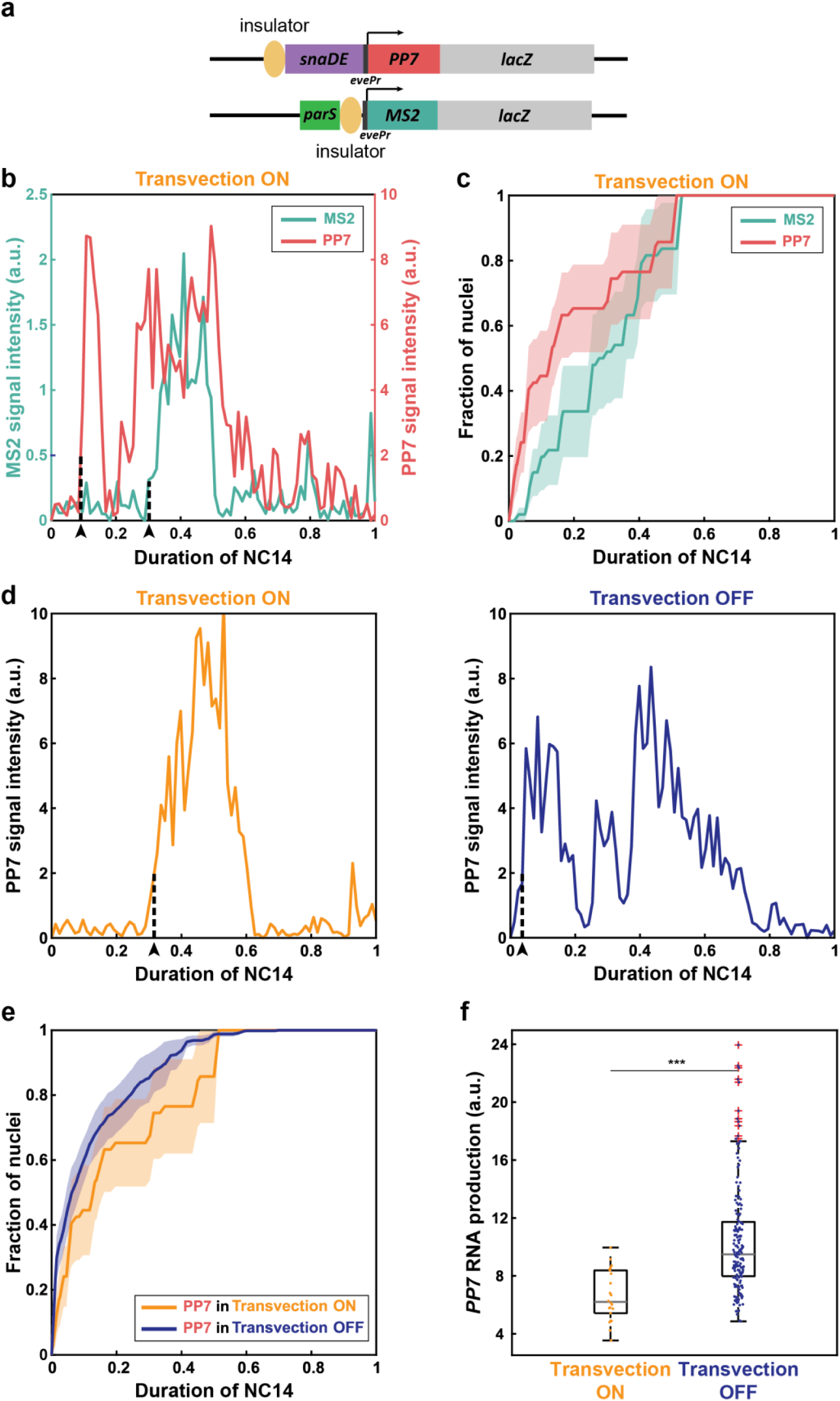
The kinetics of co-transvection reveals a competitive relationship between homologous promoters. **A)** Schematics of the DNA-labeled co-transvection assay. **B)** Representative PP7 (red) and MS2 (teal) signal trajectories of a Transvection ON nucleus. Dashed lines indicate the timing of activation of a given reporter gene. **C)** Kinetics of PP7 and MS2 signal activation of Transvection ON nuclei in NC14. The lines and shaded areas indicate the means and standard errors of PP7 (red) and MS2 (teal) signals of Transvection ON nuclei. In Transvection ON nuclei, the *cis-*linked *PP7* reporter gene has a faster activation rate than the *trans-*linked *MS2* reporter gene. 8 biological replicate embryos were analyzed. **D)** Representative PP7 signal trajectory of a Transvection ON (left) and Transvection OFF (right) nucleus. The dashed line indicates the timing of activation for the PP7 signal. **E)** The kinetics of the PP7 signal activation of Transvection ON and Transvection OFF nuclei in NC14. The lines and shaded areas indicate the mean and standard errors of Transvection ON (orange) and Transvection OFF (blue) nuclei. The *PP7* activation is delayed in the presence of *trans-*activation (Transvection ON), as opposed to in the absence of it (Transvection OFF). 8 biological replicate embryos were analyzed. **F)** The *PP7* mRNA production of Transvection ON and OFF nuclei. The *cis*-linked *PP7* reporter gene has a lower mRNA production in the presence of *trans-*activation (Transvection ON) than in the absence of it (Transvection OFF). Data of Transvection ON and OFF were collected from 22 and 183 nuclei, respectively. ***p<1E-4 using the Welch’s *t*-test.

Next, we investigated the transcriptional activity of the *cis-*linked *PP7* reporter gene when the enhancer regulates either one target promoter (in the absence of transvection) or two homologous promoters simultaneously (with transvection). Interestingly, the *cis*-linked reporter gene had a faster activation rate when *trans-*activation was absent (Fig. 4d and e). Although the enhancer preferentially interacts with the *cis*-linked promoter, the presence of the *trans-*interaction may affect the *cis*-interaction, leading to a delay in transcriptional activation in *cis*. Such delayed *cis* activation suggests that the two homologous promoters compete for the enhancer and shared transcriptional machinery when activated in the same transcription hub. Lastly, we quantified the mRNA production from the *cis*-linked *PP7* reporter gene in the absence and presence of *trans*-activation (Fig. 4f). mRNA production of the *PP7* reporter gene was reduced in transvecting nuclei, indicating that the transcriptional activity of the *MS2* reporter gene affected that of the *PP7* reporter, similar to the rate of activation. These results reveal the competitive relationship between homologous promoters and further support the existence of a transcription hub with a limiting amount of transcriptional machinery.

## Discussion

We leveraged the phenomenon of transvection to investigate the spatiotemporal activation requirements for long-distance enhancer-promoter interactions. We implemented an in *vivo* DNA-labeling technique based on the ParB/*parS* system in the co-transvection construct, where a single enhancer regulates two target promoters in *cis* and in *trans*. By tagging the *trans*-linked promoter in the enhancer-less allele with the *parS* sequence, we were able to track the dynamics of E-P interactions in living cells independent of the *trans*-transcriptional activity. This ParB/*parS* system should be applicable for other tissues and organisms to enable in *vivo* DNA visualization in real time.

Utilizing the DNA-labeled co-transvection assay that visualizes both transcription and DNA locus in living cells, we quantified the change in E-P dynamics throughout NC14, in the presence and absence of transvection. Although transient E-P proximity was observed frequently for about half the analyzed nuclei, it was insufficient to induce transcription (Fig. 1c). Since active transcription in *trans* was only observed in nuclei where the enhancer and promoter sustained *trans-*allelic proximity with an average E-P distance of 506 nm and four minutes of interactions, we conclude that persistent E-P spatial proximity is necessary and sufficient for transcription initiation. This conclusion is in line with the finding of a previous study that examined the dynamics of long-range E-P interactions in *cis.* They found that sustained E-P proximity with a mean distance of ∼350 nm is required for transcriptional activation^23,43^. Given the long distance between the enhancer and promoter at homologous positions and the low efficiency of insulator homolog pairing, the E-P distance shown with our DNA-labeled co-transvection assay could be a minimal working distance for the enhancer to interact with its promoter. This also explains the longer E-P distance during *trans-*activation, compared to the average E-P distance during *cis-*activation. Still, both the previous finding and our work indicate that active transcription is a result of sustained proximal E-P interactions, while non-active loci show more distal and transient dynamics.

Although persistent E-P proximity is shown to be important for transcription initiation in *Drosophila*, the same conclusion could not be drawn in mouse embryonic stem cells (mESCs). On the contrary, mESCs studies have found that the E-P spatial distance has no direct relationship with transcription initiation in live cells and fixed cells^24-27^. DNA FISH in mESCs even showed that E-P distance increases during active transcription^28,29^. However, most of the E-P distances in these studies fall within 500 nm regardless of the state of transcriptional activity. Interestingly, this proximity region is similar to the minimal working distance for the enhancer to activate its target promoter reported in this study. In other words, while the spatial proximity criteria may be different for varying organisms or tissue types, enhancers and their target promoters still need to be in a relative proximity of ∼500 nm to activate the target gene. A potential limitation of our *Drosophila* study arises from the enhancer’s positioning. The constant activation of the *cis*-linked reporter gene, necessary for enhancer localization, may inadvertently sequester transcription machinery and create an artificially increased distance between the *cis*- and *trans*-linked reporter genes. This scenario could influence the natural dynamics of long-distance enhancer-promoter interactions. A more comprehensive understanding of the relationship between the E-P distance and transcription requires further probing with an optimized assay with improved DNA labeling systems for both the enhancer and promoter.

The DNA-labeled co-transvection assay also elucidated the key temporal information of the initiation and termination of E-P interactions. The E-P distance dynamics revealed that the enhancer and promoter gradually approached each other through insulator-mediated pairing and remained in proximity about 4 minutes before the onset of active transcription (Fig. 3b). We hypothesize that these few minutes of E-P interactions may serve as a temporal threshold for transcriptional activation. During this period, E-P interactions could facilitate the transition from a transcriptionally "OFF" to a transcriptionally "ON" state, potentially involving *trans*-allelic proximity, recruitment of transcription machinery, or changes in chromatin accessibility^14,44-48^. Since transvection was only observed in nuclei that exhibited *trans-*allelic proximity for at least a few minutes, the ability to extend E-P interactions for around four minutes may be one of the key steps to initiate transcription and control gene expression. Furthermore, E-P proximity was maintained for around ten minutes after termination of active transcription, indicating the stability of E-P interactions and a potential primed state for rapid re-initiation of transcription or a gradual disassembly of the transcriptional complex. The E-P proximity extending from pre-initiation to post-termination of transcription resembles the conditions of a transcription hub as proposed in other studies^30,40-42^.

Through investigation of the transcriptional kinetics of *cis-* and *trans-*linked reporter genes, we have characterized a “transcription hub” environment when a single shared enhancer co-activates the target gene in *cis* and *trans*. Strikingly, transcription of the *cis*-linked reporter gene showed a delay in activation when the shared enhancer co-activated transcription at the *cis-* and *trans*-linked reporter genes (Fig. 4e). Additionally, the mRNA production at the *cis*-linked reporter gene was greater in the absence of *trans*-activation (Fig. 4f). These observations suggest an allelic competition between the homologous promoters, which supports the existence of a transcription hub with limited shared transcriptional machinery, formed through insulator-mediated homolog pairing.

However, it has yet to identify which element or elements of the transcriptional machinery are the limiting factors in the hub. Moreover, we observed multiple consecutive transcriptional bursts, instead of a long continuous transcriptional activity, despite a stable E-P association (Fig. 1d and 4b). This result implies that parameters other than the stability of the transcription hub determine the properties of transcriptional bursting.

Taken together, our high-spatiotemporal resolution live imaging study of long-range E-P interactions in *trans* offers significant insights into the mechanism of enhancer-mediated transcriptional regulation. It has enabled us to establish quantitative spatial and temporal thresholds of E-P interactions required for transcription initiation, while simultaneously elucidating the mechanistic underpinnings of the regulation of multiple promoters within a transcription hub. We established transvection as an interchromosomal mechanism to regulate gene expression in *Drosophila* and potentially in other species. Our findings not only advance the understanding of the dynamics and constraints of long-distance gene regulation, but also provide a framework for future investigations into the complex orchestration of gene expression in three-dimensional nuclear space.

## Methods

### Fly lines

The following fly lines were used in this study: *parS-gypsy-evePr-MS2-lacZ* (this study), *sp* / *cyo; Dr* / *TM3* (Bloomington Stock Center #59967), *MCP::BFP^23^*, *PCP::mKate; ParB::eGFP*^23^, *gypsy-snaDE-evePr-PP7-lacZ^30^*.

*MCP::BFP; parS-gypsy-evePr-MS2-lacZ* was generated from *MCP::BFP* and *parS-gypsy-evePr-MS2-lacZ. MCP::BFP, parS-gypsy-evePr-MS2-lacZ* line was then crossed with *PCP::mKate; ParB::eGFP* to obtain *MCP::BFP* / *PCP::mKate; ParB::eGFP* / *parS-gypsy-evePr-MS2-lacZ.* A cage of virgin female flies carrying *MCP::BFP* / *PCP::mKate; ParB::eGFP* / *parS-gypsy-evePr-MS2-lacZ* and male flies containing *gypsy-snaDE-evePr-PP7-lacZ* on the third chromosome was set up to collect the embryos carrying *parS-gypsy-evePr-MS2-lacZ* and *gypsy-snaDE-evePr-PP7-lacZ* with maternally deposited fluorescent proteins, MCP::BFP, PCP::mKate, and ParB::eGFP.

### Transgenic constructs

The *parS-gypsy-evePr-MS2-lacZ* construct was generated based on the *gypsy-evePr-MS2-lacZ* construct from Lim et al. 2018^30^. The 796 bp *parS* sequence was obtained from the Lagha Lab and amplified with primers (5’ - GCTTGCATGCCTGCAGGAC - 3’) and (5’ - GAAGTTCAAGGAATTCGC - 3’) and inserted directly upstream of the *gypsy* insulator. The plasmid was integrated into a unique landing site on the third chromosome of the *Drosophila* genome via PhiC31-mediated site-specific integration using the *VK00033* locus (Bloomington *Drosophila* Stock Center, cat#9750)^49^. BestGene Inc. generated the fly line with the transgenic construct.

### The ParB/*parS* system for in *vivo* live DNA labeling

The DNA-labeling method is derived from the parABS bacterial partitioning system that ensures proper segregation of replicons in daughter cells following division^50^. It relies on the ParB protein that binds with high specificity to a palindromic DNA sequence of 16 base pairs called *parS*. Upon binding to *parS*, the ParB dimer oligomerizes additional ParB molecules, thus nucleating a large focus of ParB proteins engaging nonspecific DNA binding in the vicinity of *parS* sites^51^. This spreading of ParB on DNA adjacent to *parS* sites allows detection of the protein focus with high signal to noise ratio (SNR) and alleviates the requirement for handling highly multimerized binding sites. Since most ParB proteins are bound through relatively weak DNA interactions, formation of the ParB/*Par* complex does not interfere with chromatin organization, transcription or accessibility. We used ParB and *parS* components from *Burkholderia cenocepacia*^52^, which do not require any additional bacterial proteins and can thus be transferred to eukaryotes, including yeast^53^, plants^54^, mammalian cells and viruses^55,56^, and *Drosophila*^23,57,58^. For locus tagging, we utilized a DNA fragment of approximately 1 kb (position 3423-4585 of *B. cenocepacia* J2315 chromosome 3)^23,53,59^, which contains two *parS* inverted repeats. The ParB2 protein sequence was C-terminally fused to eGFP and we did not introduce a nuclear localization signal. Therefore, only ParB proteins bound to *parS* remain present in the nuclear compartment, contributing to high SNR. A more extensive description of the ParB/*parS* system will be published elsewhere. The transgenic lines allowing maternal expression of ParB::eGFP (inserted at a 89B8 landing site) was a gift of Thomas Gregor^23^. All DNA constructs were made through standard cloning procedures and further detailed procedures are available upon request.

### Live Imaging

Embryos were collected from the cage after two hours at 23°C. They were dechorionated with 50 % bleach, washed, and mounted on a semipermeable membrane (Sarstedt) secured by a custom frame. Live images were taken by a Zeiss confocal microscope LSM 800 with Plan-Apochromat 40x/numerical aperture (NA) 1.3 oil-immersion objective with 50 μm pinhole and Plan-Apochromat 63x/NA 1.4 oil-immersion objective with 41 μm pinhole, and the ZEN microscopy software. Data from Figures 1, 2, 4f Supplementary Figure 1, and Supplementary Figure 2b were imaged with a 63x objective, and data from Figures 3, 4b-e, and Supplementary Figure 2c-e were imaged with a 40x objective. All images were in 16 bits and contained 512 x 512 pixels. The 40x images had a 1.5x zoom and a pixel size of 208 nm and over 17 Z-stack slices with a 0.67 μm interval. The 63x images had a 1.3x zoom and a pixel size of 152 nm and over 32 Z-stack slices with a 0.34 μm interval. The temporal resolutions were 35.39 s for 40x images and 36.69 s for 63x images. Three lasers, 405 nm, 488 nm, and 561 nm, were used to visualize MCP::BFP, ParB::eGFP, and PCP::mKate, respectively.

### Image Analysis

All live images were compiled in ImageJ and analyzed in MATLAB (R2018b, MathWorks). The three-dimensional raw images were compiled into two-dimensional movies through a maximal Z projection in ImageJ. From the onset of NC14 to gastrulation, nuclei were segmented by masks via a custom MATLAB code and then manually corrected in ImageJ. To extract PP7 and MS2 signals from each nucleus, the two highest pixel values within each nucleus mask at each time frame were located and the averages of their eight neighboring pixel values were calculated for each of the two highest-intensity pixel positions. The pixel value with the higher neighboring average was denoted as the fluorescent signal intensity (PP7 or MS2) of the nucleus at a given time frame. The x and y coordinates of each denoted fluorescent signal of each nucleus were extracted over time.

### Enhancer-promoter Distance Calculation

The three-dimensional enhancer-promoter distances were calculated as 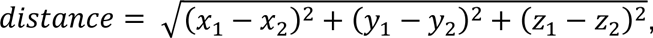 using the x-, y-, and z-coordinates of each MS2, PP7, or *parS* signal. The x- and y-coordinates of each signal were extracted from the two-dimensional max-projected live images. The z-coordinate of each signal was extracted from the three-dimensional raw live images. The z-signal trajectory at the x- and y-coordinates of the fluorescent signals was extracted at each time frame. The z-coordinate was approximated as the peak of the one-term Gaussian approximation of the z-signal trajectory. The z-coordinates were manually selected as one of the peaks with two- or three-term Gaussian approximation when the one-term Gaussian approximation did not match the signal trajectories over Z-stack.

## Acknowledgements

We thank members of the Lim lab for helpful discussions and comments on the manuscript. We are grateful to Kerstin Bystricky and Jean-Yves Bouet for their help in implementing the ParB/parS system in flies, Mounia Lagha for providing the parS-containing plasmid, Thomas Gregor for providing the MCP::BFP and PCP::mKate; ParB::eGFP fly stocks. This work was supported by the National Science Foundation CAREER MCB 2044613 awarded to B.L., as well as by ANR (Chrononet) and Fondation pour la Recherche Médicale (DEQ20170336739) to F.P..

## Author Contribution

H.D. and B.L. conceived the project and designed the experiments. H.D. performed all experiments and analyzed data. P.V. and F.P. implemented the ParB/ParS system. B.L. supervised the project. H.D., F.P., and B.L. wrote the manuscript, and P.V. provided edits.

## Reference

1 Claringbould, A. & Zaugg, J. B. Enhancers in disease: molecular basis and emerging treatment strategies. Trends Mol Med 27, 1060–1073 (2021). 10.1016/j.molmed.2021.07.012

2 Downes, D. J. & Hughes, J. R. Natural and Experimental Rewiring of Gene Regulatory Regions. Annu Rev Genomics Hum Genet 23, 73–97 (2022). 10.1146/annurev-genom-112921-010715

3 Carroll, S. B. Evo-devo and an expanding evolutionary synthesis: a genetic theory of morphological evolution. Cell 134, 25–36 (2008). 10.1016/j.cell.2008.06.030

4 Hill, M. S., Vande Zande, P. & Wittkopp, P. J. Molecular and evolutionary processes generating variation in gene expression. Nat Rev Genet 22, 203–215 (2021). 10.1038/s41576-020-00304-w

5 Hnisz, D. et al. Super-enhancers in the control of cell identity and disease. Cell 155, 934–947 (2013). 10.1016/j.cell.2013.09.053

6 Miguel-Escalada, I., Pasquali, L. & Ferrer, J. Transcriptional enhancers: functional insights and role in human disease. Curr Opin Genet Dev 33, 71–76 (2015). 10.1016/j.gde.2015.08.009

7 Yang, J. H. & Hansen, A. S. Enhancer selectivity in space and time: from enhancer-promoter interactions to promoter activation. Nat Rev Mol Cell Biol 25, 574–591 (2024). 10.1038/s41580-024-00710-6

8 Levine, M. & Tjian, R. Transcription regulation and animal diversity. Nature 424, 147–151 (2003). 10.1038/nature01763

9 Levine, M., Cattoglio, C. & Tjian, R. Looping back to leap forward: transcription enters a new era. Cell 157, 13–25 (2014). 10.1016/j.cell.2014.02.009

10 Muller, M. M., Gerster, T. & Schaffner, W. Enhancer sequences and the regulation of gene transcription. Eur J Biochem 176, 485–495 (1988). 10.1111/j.1432-1033.1988.tb14306.x

11 Schaffner, W. Enhancers, enhancers - from their discovery to today’s universe of transcription enhancers. Biol Chem 396, 311–327 (2015). 10.1515/hsz-2014-0303

12 Maniatis, T., Goodbourn, S. & Fischer, J. A. Regulation of inducible and tissue-specific gene expression. Science 236, 1237–1245 (1987). 10.1126/science.3296191

13 Kvon, E. Z. et al. Genome-scale functional characterization of Drosophila developmental enhancers in vivo. Nature 512, 91–95 (2014). 10.1038/nature13395

14 Ghavi-Helm, Y. et al. Enhancer loops appear stable during development and are associated with paused polymerase. Nature 512, 96–100 (2014). 10.1038/nature13417

15 Sanyal, A., Lajoie, B. R., Jain, G. & Dekker, J. The long-range interaction landscape of gene promoters. Nature 489, 109–113 (2012). 10.1038/nature11279

16 Long, H. K., Prescott, S. L. & Wysocka, J. Ever-Changing Landscapes: Transcriptional Enhancers in Development and Evolution. Cell 167, 1170–1187 (2016). 10.1016/j.cell.2016.09.018

17 Furlong, E. E. M. & Levine, M. Developmental enhancers and chromosome topology. Science 361, 1341–1345 (2018). 10.1126/science.aau0320

18 Chathoth, K. T. et al. The role of insulators and transcription in 3D chromatin organization of flies. Genome Res 32, 682–698 (2022). 10.1101/gr.275809.121

19 Calhoun, V. C. & Levine, M. Long-range enhancer-promoter interactions in the Scr-Antp interval of the Drosophila Antennapedia complex. Proc Natl Acad Sci U S A 100, 9878–9883 (2003). 10.1073/pnas.1233791100

20 Banerji, J., Olson, L. & Schaffner, W. A lymphocyte-specific cellular enhancer is located downstream of the joining region in immunoglobulin heavy chain genes. Cell 33, 729–740 (1983). 10.1016/0092-8674(83)90015-6

21 Gillies, S. D., Morrison, S. L., Oi, V. T. & Tonegawa, S. A tissue-specific transcription enhancer element is located in the major intron of a rearranged immunoglobulin heavy chain gene. Cell 33, 717–728 (1983). 10.1016/0092-8674(83)90014-4

22 Mercola, M., Wang, X. F., Olsen, J. & Calame, K. Transcriptional enhancer elements in the mouse immunoglobulin heavy chain locus. Science 221, 663–665 (1983). 10.1126/science.6306772

23 Chen, H. et al. Dynamic interplay between enhancer-promoter topology and gene activity. Nat Genet 50, 1296–1303 (2018). 10.1038/s41588-018-0175-z

24 Alexander, J. M. et al. Live-cell imaging reveals enhancer-dependent Sox2 transcription in the absence of enhancer proximity. Elife 8 (2019). 10.7554/eLife.41769

25 Sexton, T. et al. Competition between transcription and loop extrusion modulates promoter and enhancer dynamics. Res Sq (2023). 10.21203/rs.3.rs-3164817/v1

26 Taylor, T. et al. Transcriptional regulation and chromatin architecture maintenance are decoupled functions at the Sox2 locus. Genes Dev 36, 699–717 (2022). 10.1101/gad.349489.122

27 Chakraborty, S. et al. Enhancer-promoter interactions can bypass CTCF-mediated boundaries and contribute to phenotypic robustness. Nat Genet 55, 280–290 (2023). 10.1038/s41588-022-01295-6

28 Benabdallah, N. S. et al. Decreased Enhancer-Promoter Proximity Accompanying Enhancer Activation. Mol Cell 76, 473–484 e477 (2019). 10.1016/j.molcel.2019.07.038

29 Kane, L. et al. Cohesin is required for long-range enhancer action at the Shh locus. Nat Struct Mol Biol 29, 891–897 (2022). 10.1038/s41594-022-00821-8

30 Lim, B., Heist, T., Levine, M. & Fukaya, T. Visualization of Transvection in Living Drosophila Embryos. Mol Cell 70, 287–296 e286 (2018). 10.1016/j.molcel.2018.02.029

31 Lewis, E. B. The Theory and Application of a New Method of Detecting Chromosomal Rearrangements in Drosophila-Melanogaster. Am Nat 88, 225–239 (1954). Doi 10.1086/281833

32 Peifer, M. & Bender, W. The anterobithorax and bithorax mutations of the bithorax complex. EMBO J 5, 2293–2303 (1986). 10.1002/j.1460-2075.1986.tb04497.x

33 Duncan, I. W. Transvection effects in Drosophila. Annu Rev Genet 36, 521–556 (2002). 10.1146/annurev.genet.36.060402.100441

34 Harrison, D. A., Gdula, D. A., Coyne, R. S. & Corces, V. G. A leucine zipper domain of the suppressor of Hairy-wing protein mediates its repressive effect on enhancer function. Genes Dev 7, 1966–1978 (1993). 10.1101/gad.7.10.1966

35 Kravchenko, E. et al. Pairing between gypsy insulators facilitates the enhancer action in trans throughout the Drosophila genome. Mol Cell Biol 25, 9283–9291 (2005). 10.1128/MCB.25.21.9283-9291.2005

36 Bertrand, E. et al. Localization of ASH1 mRNA particles in living yeast. Mol Cell 2, 437–445 (1998). 10.1016/s1097-2765(00)80143-4

37 Larson, D. R., Zenklusen, D., Wu, B., Chao, J. A. & Singer, R. H. Real-time observation of transcription initiation and elongation on an endogenous yeast gene. Science 332, 475–478 (2011). 10.1126/science.1202142

38 Garcia, H. G., Tikhonov, M., Lin, A. & Gregor, T. Quantitative imaging of transcription in living Drosophila embryos links polymerase activity to patterning. Curr Biol 23, 2140–2145 (2013). 10.1016/j.cub.2013.08.054

39 Negre, N. et al. A comprehensive map of insulator elements for the Drosophila genome. Plos Genet 6, e1000814 (2010). 10.1371/journal.pgen.1000814

40 Fukaya, T., Lim, B. & Levine, M. Enhancer Control of Transcriptional Bursting. Cell 166, 358–368 (2016). 10.1016/j.cell.2016.05.025

41 Hnisz, D., Shrinivas, K., Young, R. A., Chakraborty, A. K. & Sharp, P. A. A Phase Separation Model for Transcriptional Control. Cell 169, 13–23 (2017). 10.1016/j.cell.2017.02.007

42 Liu, Z. & Tjian, R. Visualizing transcription factor dynamics in living cells. J Cell Biol 217, 1181–1191 (2018). 10.1083/jcb.201710038

43 Bruckner, D. B., Chen, H., Barinov, L., Zoller, B. & Gregor, T. Stochastic motion and transcriptional dynamics of pairs of distal DNA loci on a compacted chromosome. Science 380, 1357–1362 (2023). 10.1126/science.adf5568

44 Dixon, J. R. et al. Chromatin architecture reorganization during stem cell differentiation. Nature 518, 331–336 (2015). 10.1038/nature14222

45 Krijger, P. H. et al. Cell-of-Origin-Specific 3D Genome Structure Acquired during Somatic Cell Reprogramming. Cell Stem Cell 18, 597–610 (2016). 10.1016/j.stem.2016.01.007

46 Raser, J. M. & O’Shea, E. K. Control of stochasticity in eukaryotic gene expression. Science 304, 1811–1814 (2004). 10.1126/science.1098641

47 Voss, T. C. & Hager, G. L. Dynamic regulation of transcriptional states by chromatin and transcription factors. Nat Rev Genet 15, 69–81 (2014). 10.1038/nrg3623

48 Sanchez, A., Garcia, H. G., Jones, D., Phillips, R. & Kondev, J. Effect of promoter architecture on the cell-to-cell variability in gene expression. PLoS Comput Biol 7, e1001100 (2011). 10.1371/journal.pcbi.1001100

49 Venken, K. J., He, Y., Hoskins, R. A. & Bellen, H. J. P[acman]: a BAC transgenic platform for targeted insertion of large DNA fragments in D. melanogaster. Science 314, 1747–1751 (2006). 10.1126/science.1134426

50 Surtees, J. A. & Funnell, B. E. Plasmid and chromosome traffic control: how ParA and ParB drive partition. Curr Top Dev Biol 56, 145–180 (2003). 10.1016/s0070-2153(03)01010-x

51 Sanchez, A. et al. Stochastic Self-Assembly of ParB Proteins Builds the Bacterial DNA Segregation Apparatus. Cell Syst 1, 163–173 (2015). 10.1016/j.cels.2015.07.013

52 Dubarry, N., Pasta, F. & Lane, D. ParABS systems of the four replicons of Burkholderia cenocepacia: new chromosome centromeres confer partition specificity. J Bacteriol 188, 1489–1496 (2006). 10.1128/JB.188.4.1489-1496.2006

53 Saad, H. et al. DNA dynamics during early double-strand break processing revealed by non-intrusive imaging of living cells. Plos Genet 10, e1004187 (2014). 10.1371/journal.pgen.1004187

54 Meschichi, A. et al. ANCHOR: A Technical Approach to Monitor Single-Copy Locus Localization in Planta. Front Plant Sci 12, 677849 (2021). 10.3389/fpls.2021.677849

55 Mariame, B. et al. Real-Time Visualization and Quantification of Human Cytomegalovirus Replication in Living Cells Using the ANCHOR DNA Labeling Technology. J Virol 92 (2018). 10.1128/JVI.00571-18

56 Germier, T. et al. Real-Time Imaging of a Single Gene Reveals Transcription-Initiated Local Confinement. Biophys J 113, 1383–1394 (2017). 10.1016/j.bpj.2017.08.014

57 Gomez-Lamarca, M. J. et al. Activation of the Notch Signaling Pathway In Vivo Elicits Changes in CSL Nuclear Dynamics. Dev Cell 44, 611–623 e617 (2018). 10.1016/j.devcel.2018.01.020

58 Delker, R. K., Munce, R. H., Hu, M. & Mann, R. S. Fluorescent labeling of genomic loci in Drosophila imaginal discs with heterologous DNA-binding proteins. Cell Rep Methods 2, 100175 (2022). 10.1016/j.crmeth.2022.100175

59 Germier, T., Audibert, S., Kocanova, S., Lane, D. & Bystricky, K. Real-time imaging of specific genomic loci in eukaryotic cells using the ANCHOR DNA labelling system. Methods 142, 16–23 (2018). 10.1016/j.ymeth.2018.04.008

